# Bacterial Virulence Genes Detected by Metagenomic Sequencing in the Cystic Fibrosis Airway Microbiome

**DOI:** 10.64898/2026.05.19.726200

**Authors:** Medha L. Valluri, Brennan Harmon, Aszia Burrell, Andrea Hahn

## Abstract

**Background:** Cystic fibrosis (CF) is an autosomal recessive genetic disorder that leads to chronic infection and mucus retention in the lungs, with lung function gradually deteriorating through recurrent pulmonary exacerbations (PEx). Virulence factors (VFs) of *Pseudomonas aeruginosa* and *Staphylococcus aureus* are thought to contribute to pulmonary exacerbations. Our study objective was to identify VF genes related to PEx, high *Pseudomonas* abundance, and high *Staphylococcus* abundance in persons with CF (pwCF).

**Methods:** This was an ancillary study of pwCF treated with IV antibiotics for PEx between 2016-2020 at Children’s National Hospital. Using shotgun metagenomics and ShortBRED, we identified bacterial VF genes and used DESeq2 to determine differential expression of VF genes across comparators.

**Results:** Twenty-two PwCF experienced 43 PEx. The study cohort had a mean age of 14.6 years, 41% female, 59% white, 36% Hispanic, and 45% had an F508del homozygous CFTR mutation. Minimal differences in VF gene abundance were identified across clinical state. The most differentially increased VF genes found in *Pseudomonas* high samples were associated with an aminotransferase (log2FC 25.9), flagellar biosynthesis (log2FC 8.3), and type VI secretion systems (log2FC 8.2). The most differentially increased VF genes found in *Staphylococcus* high samples were an exotoxin (log2FC 26.7), macrolide phosphotransferase (log2FC 25.8), pathogenicity island proteins (log2FC 25.2 and 24.7), and VOC family proteins (log2FC 24.8).

**Conclusions:** These findings demonstrate that specific VFs associated with immune modulation, motility secretion systems, bacterial motility, and antibiotic resistance are related to *P. aeruginosa* and *S. aureus* abundance, providing potential targets for more personalized antimicrobial interventions.

## Introduction

Cystic fibrosis (CF) is an autosomal recessive genetic disorder that leads to chronic infection and mucus retention in the lungs (1). According to Flume et al., lung function for persons with CF (pwCF) gradually deteriorates with periodic sudden intensifications of breathing-related symptoms, commonly called pulmonary exacerbations (2). The clinical signs of a pulmonary exacerbation may include heightened coughing, greater mucus production, difficulty breathing, chest discomfort, decreased appetite, weight loss, and reduced lung function. Two common infections during pulmonary exacerbations for pwCF are *Staphylococcus aureus* and *Pseudomonas aeruginosa* (3). According to Israyilova et al., *S. aureus* continues to be the bacterial infection most contracted by young children with CF within their initial year of life (4). While the prevalence of *P. aeruginosa* and *S. aureus* has declined over time, both remain the most common respiratory microorganisms in pwCF (5).

Virulence factors (VFs), alternatively termed effectors, are substances such as molecules, proteins, or toxins that organisms employ to adhere to or penetrate host cells, suppress or escape immune defenses, or trigger cellular death (6,7). For *P. aeruginosa*, one of these VFs is TseT, which is upregulated in response to antiviral signaling in a host (8). This ensures that *P. aeruginosa* remains prevalent in the environment and ultimately lowers microbial diversity and lung function in CF airways. Additionally, *P. aeruginosa* ensures survival through its Fur protein, which activates when the host system has low iron values (9). Fur participates in generating toxins, creating more developed biofilms, and facilitating quorum sensing*. S. aureus*, the other most common bacteria in pwCF, can also adapt to host environments to ensure survival. For example, *S. aureus* has several virulence mechanisms that allow it to adapt to the CF lung, including biofilm formation, misdirection of the immune system through secretion of toxins, and the ability to develop small-colony variants (10).

Pulmonary exacerbations (PEx) are typically treated with a combination of airway clearance and antibiotics such as aminoglycosides or β-lactams for varying lengths of time (2,11). The conventional treatment strategy for *P. aeruginosa* infection associated with PEx involves administering multiple antibiotics simultaneously, with physicians generally choosing antimicrobials based on pathogen susceptibility profiles (12). However, for pwCF, identifying antibiotics effective against all detected pathogens may not be achievable. *P. aeruginosa* can adapt its VF production during chronic infection, which has frequently been shown to occur in the CF lung (13). Additionally, according to Buckley et al., *S. aureus* makes several virulence determinants, which interfere with current and experimental treatment options (14). As mentioned above, *S. aureus* and *P. aeruginosa* both stimulate increased biofilm production (10), which is detrimental towards existing paths of treatment for pulmonary exacerbations. Biofilms, which are a dense cluster of microbial cells, increase antibiotic resistance by creating a protective layer around bacteria, blocking the entry of antibiotics (15). This leaves a gap in effective treatment options for pwCF.

Taken together, we hypothesize that understanding changes in bacterial VFs around the time of PEx may provide insights into more effective antibiotic treatment options. To test our hypothesis, we will determine whether bacterial VFs detected by shotgun metagenomics in CF sputum are differentially abundant based on clinical state or dominant pathogen. In this ancillary study, our primary aim is to compare bacterial VF abundance across three clinical states: PEx, completion of antibiotic treatment, and follow-up (16). Our secondary aim is to compare the relationship between the relative abundance of *P. aeruginosa* and *S. aureus* and VF abundance. By tracking VF dynamics across these three clinical timepoints, we aim to identify specific bacterial traits that may contribute to PEx. We will also identify VFs in our dataset specifically related to the two most common bacterial infections in pwCF. This knowledge could lead to more targeted antimicrobial strategies, earlier intervention protocols, and personalized treatment regimens based on the virulence profile of an individual’s respiratory microbiome.

## Materials and Methods

### Study Design

This ancillary study uses data collected from a prior observational study of pwCF who received intravenous (IV) antibiotics for treatment of PEx (16). Study participants were observed at Children’s National Hospital in Washington, DC from 2016 to 2020. The study received institutional review board approval from Children’s National Hospital under two protocols (Pro6781, 8 Dec 2015, and Pro10528, 31 Aug 2018). Adult participants aged 18 and older gave written informed consent, while parental consent was secured for minors under 18. Children ages 11-17 also provided written assent. The study followed the Declaration of Helsinki. Clinical data and respiratory samples were collected at three time points: hospital admission for PEx (E), completion of antibiotic therapy (T), and the subsequent follow-up appointment (F), provided no additional antibiotics were given between treatment completion and the follow-up time point. In order to be eligible for the study, the respiratory specimen collected at PEx had to be gathered from the lower respiratory tract, such as sputum or bronchoalveolar lavage (BAL). Demographic data gathered included age, weight, height, race, ethnicity, and CFTR genotype.

### Sample Collection and Processing

As part of our larger biobank study, sputum and BAL samples were collected in sterile containers and oropharyngeal (OP) swabs were collected using Eswab with Amies transport medium (Copan). All specimens were maintained at 4°C prior to processing. For homogenization, sputum and BAL samples were combined with equal volumes of Sputasol (ThermoFisher) and sterile normal saline, then vortexed and incubated at 37°C in a heated bead bath for 10 minutes. The homogenized sputum and BAL samples, along with the Amies medium from the OP swabs, were then transferred into microcentrifuge tubes and centrifuged at 12,000*g* for 10 minutes to pellet cellular material. Following centrifugation, the supernatants were separated from the pellets, and both fractions were stored independently at −80°C.

### DNA Extraction and Metagenomic Sequencing

Frozen cellular pellets were thawed and resuspended in sterile phosphate buffered saline (PBS) before DNA extraction. DNA was extracted using a QIAamp DNA Microbiome Kit (Qiagen) according to the manufacturer’s instructions. DNA concentration and integrity were assessed using Qubit (Thermofisher Scientific) and Bioanalyzer (Agilent), respectively. Sequencing libraries were prepared with the Nextera XT Library Prep Kit (Illumina), and 23-30 libraries per sequencing run were processed on an Illumina NextSeq 500 platform using a Mid-Output 2×150 cycle kit. Sequencing yielded an average of 5.8 million reads per sample (range: 670,000 to 21 million). After removing residual human sequences using KneadData (17), an average of 1.6 million reads per sample remained (range: 38,000 to 8 million).

### Bioinformatic and Statistical Analyses

Bacterial sequencing data was analyzed using ShortBRED, a program designed to identify protein families of interest from shotgun metagenomic sequencing data (18). VF gene counts were imported into Rstudio v4.4.1 (19), and packages *DESeq2* v1.46.0 (20), *tidyverse* v2.0.0 (21), *phyloseq* v1.50.0 (22), and *dplyr* v1.1.4 (23) were used to analyze differences in the VF gene abundance between the three clinical states (i.e., E, T, and F) and based on *S. aureus* and *P. aeruginosa* relative abundance which had previously been determined for this dataset using PathoScope 2.0 (16,24). Samples with a relative abundance of *S. aureus* and *P. aeruginosa* above 10% were considered high, while samples with a relative abundance of *S. aureus* and *P. aeruginosa* below 10% were considered low. Bacterial VFs found to be significantly different using *DESeq2* were then investigated in the reference marker database (i.e., ShortBRED_VF_2017_markers, available for download at https://huttenhower.sph.harvard.edu/shortbred/) to collect the protein sequences. The protein sequences were then searched for in pBLAST to find the VF name reported in our results, using default search settings.

## Results

### Study Cohort Demographics and Clinical Characteristics

Twenty-two participants with CF were enrolled in this study. Participant demographics and clinical characteristics are summarized in Table 1. Of the 22 participants, 9 were female. The cohort was racially diverse, with 13 participants identifying as White, 8 as Other Race, and 1 as Black. Eight participants identified as Hispanic. The mean age at first PEx and study enrollment was 14.6 years (σ = 5.0).

**Table 1.**
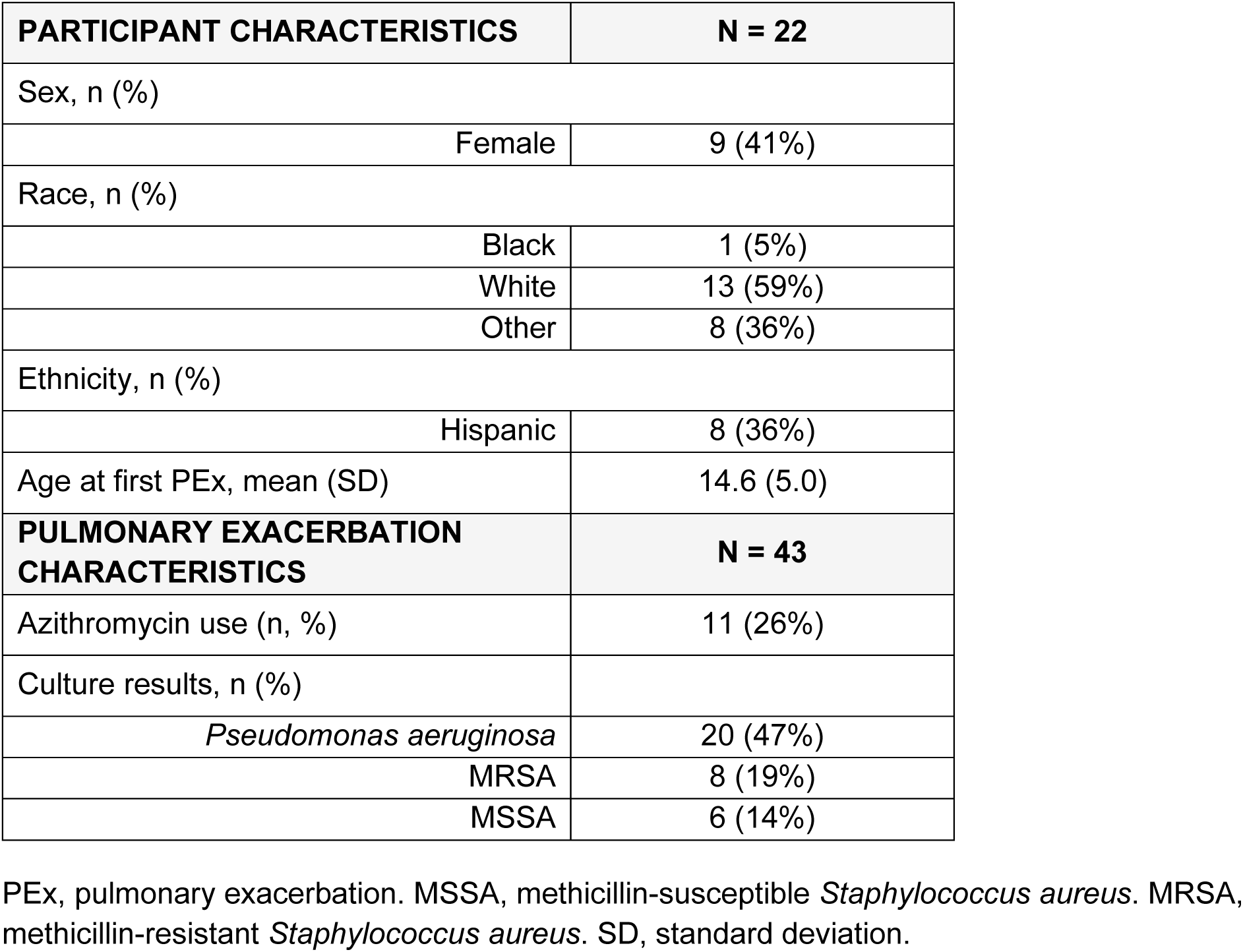
Study Participant Characteristics.

| **PARTICIPANT CHARACTERISTICS** | **N = 22** |
| --- | --- |
| Sex, n (%) |  |
| Female | 9 (41%) |
| Race, n (%) |  |
| Black | 1 (5%) |
| White | 13 (59%) |
| Other | 8 (36%) |
| Ethnicity, n (%) |  |
| Hispanic | 8 (36%) |
| Age at first PEx, mean (SD) | 14.6 (5.0) |
| **PULMONARY EXACERBATION CHARACTERISTICS** | **N = 43** |
| Azithromycin use (n, %) | 11 (26%) |
| Culture results, n (%) |  |
| *Pseudomonas aeruginosa* | 20 (47%) |
| MRSA | 8 (19%) |
| MSSA | 6 (14%) |

PEx, pulmonary exacerbation. MSSA, methicillin-susceptible *Staphylococcus aureus*. MRSA, methicillin-resistant *Staphylococcus aureus*. SD, standard deviation.

The 22 study participants experienced 43 PEx during the study period. Concurrent airway culture results at the time of PEx indicated that *P. aeruginosa* was the most prevalent organism, identified in 20 PEx, followed by methicillin-resistant *S. aureus* (MRSA) in 8 PEx and methicillin-susceptible *S. aureus* (MSSA) in 6 PEx. Respiratory samples were collected at each PEx timepoint (E); however, 10 completion of antibiotic treatment samples (T) and 6 follow-up samples (F) were not collected. A total of 112 samples successfully passed quality control and underwent shotgun metagenomic sequencing. Of these 112 metagenomic sequencing data files analyzed with ShortBRED, 75 had VF genes detected and were used in our subsequent analyses.

### Differential Abundance of Virulence Factors by Clinical State

A differential abundance analysis of VF genes was conducted between the three clinical state groups (E versus T versus F, Supplemental Figure 1). Minimal differences in VF gene abundance were noted based on clinical state (FDR < 0.05). When comparing PEx versus completion of antibiotic treatment, only two VF genes were significantly increased in PEx samples while one VF gene was increased in the completion of antibiotic treatment samples (Figure 1A). Likewise, when comparing PEx versus follow-up, only two VF genes were increased in PEx samples while two VF genes were increased in follow-up samples (Figure 1B). These findings suggest that the overall VF gene repertoire captured in metagenomic samples was not significantly altered by acute exacerbation status in this cohort.

**Figure 1.**
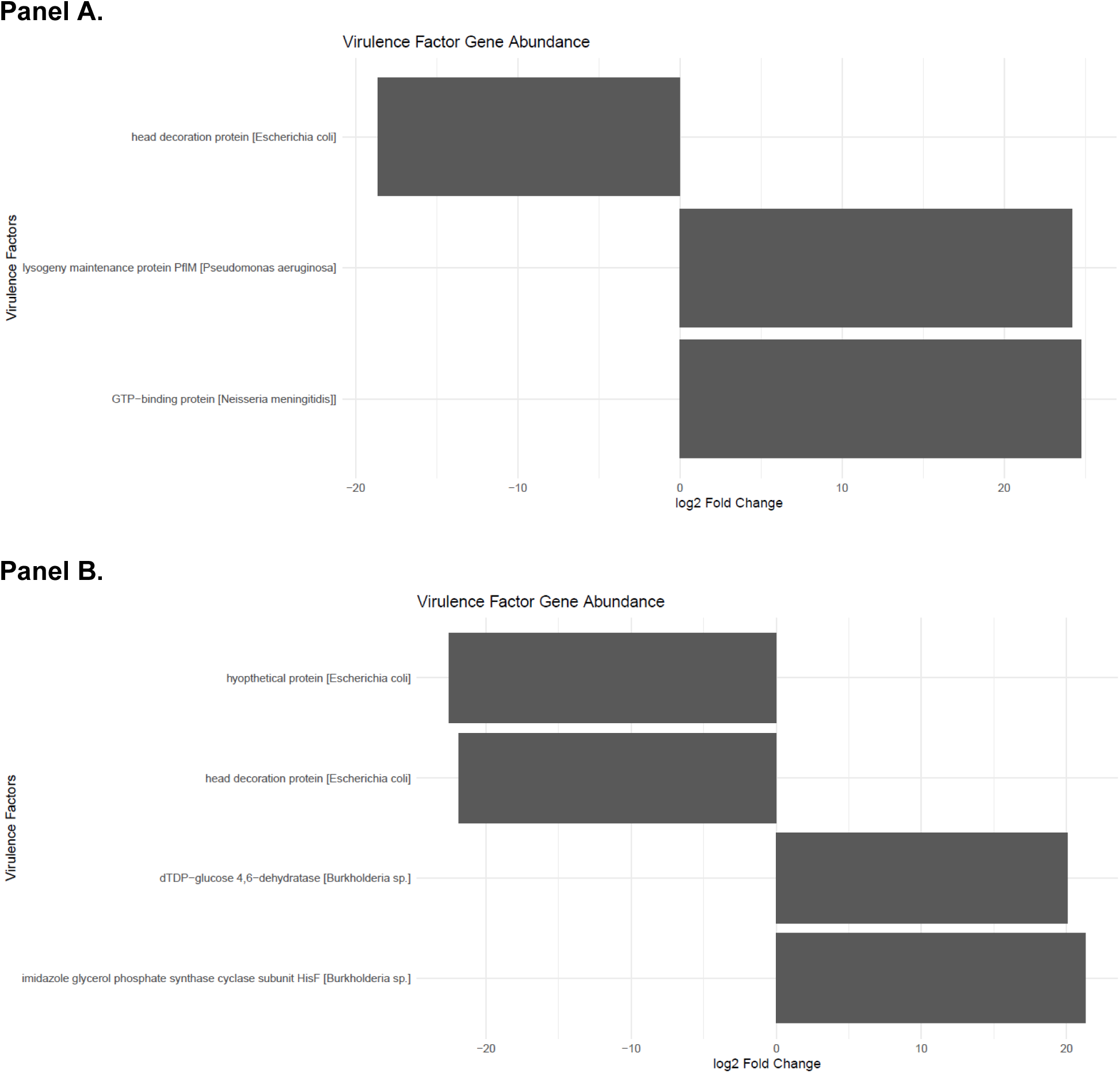
Panel A. Virulence factor gene log2 fold change between pulmonary exacerbation and completion of antibiotic treatment. Positive log2 fold change indicates higher gene abundance in the pulmonary exacerbation samples, while negative log2 fold change indicates higher gene abundance in the completion of antibiotic treatment samples. **Panel B. Virulence factor gene log2 fold change between pulmonary exacerbation and follow-up.** Positive log2 fold change indicates higher gene abundance in the pulmonary exacerbation samples, while negative log2 fold change indicates higher gene abundance in the follow-up samples.

### Differential Virulence Factor Abundance by Pseudomonas aeruginosa and Staphylococcus aureus Relative Abundance

To further characterize the virulence potential of the dominant airway pathogens, samples were stratified by relative abundance of *P. aeruginosa* and *S. aureus* into high- and low-abundance groups, and differential abundance analyses of VF genes were performed within each pathogen group.

### Pseudomonas aeruginosa Virulence Factors

Among samples stratified by *P. aeruginosa* abundance (Supplemental Figure 2), 391 VF genes were differentially increased in high-*Pseudomonas* samples, whereas 190 VF genes were differentially decreased in high-*Pseudomonas* samples (FDR < 0.05). The VF genes with the highest increase in the high-*Pseudomonas* group are shown in Figure 2. Aminotransferase degT was the most differentially abundant, with a log2FC of 25.9. Flagellar biosynthesis protein fliQ, a component of the bacterial flagellar assembly machinery associated with bacterial motility, was the next most abundant VF genes in high-*Pseudomonas* samples with a log2FC of 8.3. Other VF genes that were differentially abundant and associated with motility included type 4a pilus minor pilin pilX (log2FC 7.9), type IV pilus secretin pilQ (log2FC 7.5), and flagellar basal body rod protein flgB (log2FC 7.3). The type VI secretion system (T6SS) baseplate subunit tssG was the third most differentially abundant VF (log2FC 8.2). Other VF genes associated with bacterial secretion systems increased in high-*Pseudomonas* samples included type IV secretion system (T4SS) baseplate subunit tssF (log2FC 7.6), SycD/LcrH family type III secretion system (T3SS) chaperone pcrH (log2FC 7.5), type II secretion system (T2SS) F family protein (log2FC 7.3), and T3SS effector bifunctional cytotoxin exoenzyme S (log2 FC 7.1).

**Figure 2.**
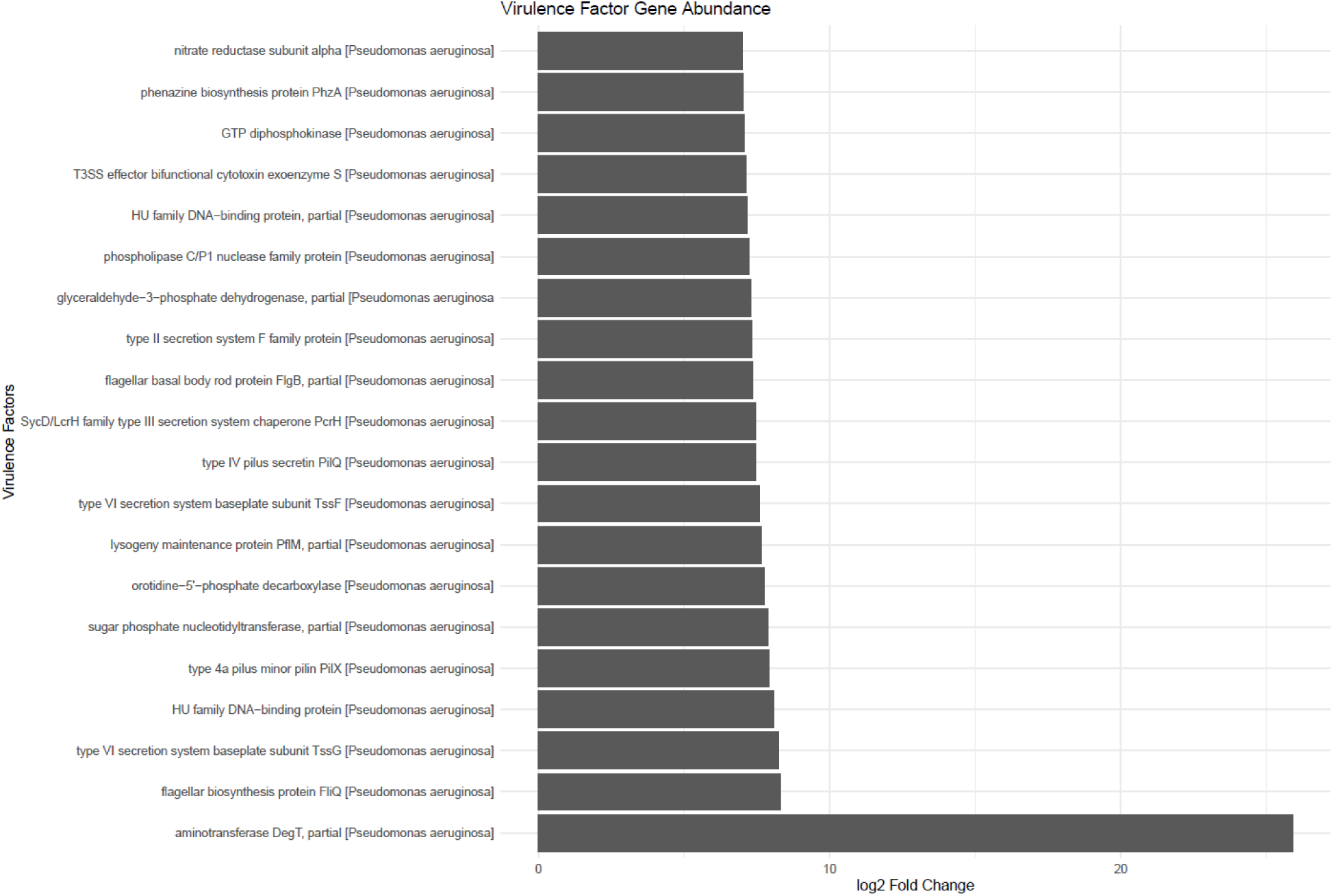
*Pseudomonas aeruginosa* virulence factor gene log2 fold change between high and low *Pseudomonas* abundance groups. Only the top 22 virulence-associated genes are shown. Positive log2 fold change indicates higher gene abundance in the high-*Pseudomonas* group.

### Staphylococcus aureus Virulence Factors

Differential abundance analysis of *S. aureus* VF genes between high- and low-*Staphylococcus* abundance groups (Supplemental Figure 3) found that 195 VF genes were differentially increased in high-*Staphylococcus* samples, whereas 491 VF genes were differentially decreased in high-*Staphylococcus* samples (FDR < 0.05). This distinct set of enriched VF genes in the high-*Staphylococcus* group are shown in Figure 3. Exotoxin beta-grasp domain-containing protein was the most differentially increased VF with a log2FC of 26.7. Other exotoxins significantly increased in high-*Staphylococcus* samples included superantigen-like protein SSL9 and superantigen-like protein SSL11 (each with a log2FC of 7.2). Additionally, pathogenicity island proteins (1) and (2) were among the most highly differentially abundant genes (log2FC 25.2 and 24.7, respectively), and they also carry genes encoding several superantigen toxins. Two VF genes associated with antibiotic resistance were also among the most differentially abundant genes: Mph(C) family macrolide 2′-phosphotransferase (log2FC 25.8) and tetracycline resistance protein (log2FC 10.7). Lastly, VOC family proteins were also enriched in high-*Staphylococcus* samples (log2FC 24.8).

**Figure 3.**
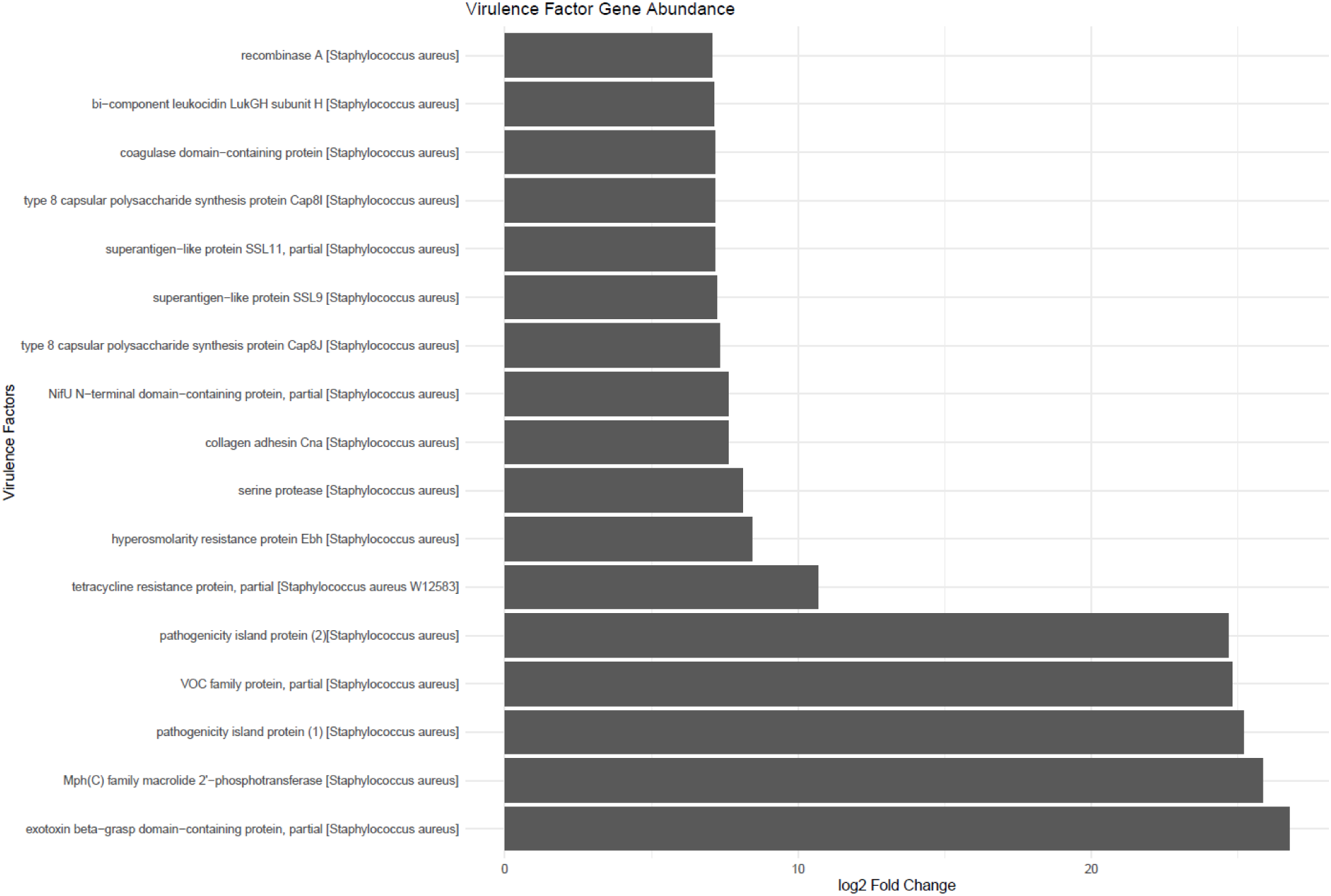
*Staphylococcus aureus* virulence factor gene log2 fold change between high and low *Staphylococcus* abundance groups. Only virulence-associated genes are shown with log2 fold change values greater than seven were included. Positive log2 fold change indicates higher gene abundance in the high-*Staphylococcus* group.

## Discussion

This study employed bioinformatic analysis to characterize VF gene abundance in the airways of pwCF. Three primary comparisons were examined: differential VF gene abundance by clinical state, differential abundance by high vs. low *P. aeruginosa* burden, and differential abundance by high vs. low *S. aureus* burden. The cohort was predominantly colonized by *P. aeruginosa* (47%), with substantial proportions of MRSA (19%) and MSSA (14%), consistent with the expected microbiological profile of a pediatric and young adult CF population.

### Virulence Factor Abundance and Clinical State

The limited number of statistically significant differences in VF gene abundance between clinical states was somewhat unexpected. The hypothesis that exacerbation would be associated with a measurable increase in VF gene abundance, driven by acute pathogen activity, was not supported by these data. Though we did find a few significant differences, they were not consistently associated with a specific bacterial pathogen being increased in PEx samples across both comparisons (E vs T and E vs F). Although *Escherichia coli* head decoration protein was decreased in PEx samples in both comparisons, this protein is likely related to an *E. coli* bacteriophage (25) and so unlikely to be a major contributor to PEx onset. While this result may initially appear counterintuitive, it is consistent with an emerging body of literature suggesting that onset PEx is not directly associated with pathogen burden. *P. aeruginosa* absolute and relative abundance are frequently found to be at similar levels before, during, and after exacerbation, particularly in older patients (26,27), and repeat PEx are most often associated with the presence of a similar strain (28). This suggests that acute exacerbations may not be uniformly driven by sudden increases in pathogen burden or VF gene expression.

An additional consideration is that *P. aeruginosa* undergoes substantial adaptive evolution during chronic CF infection, including the progressive downregulation of many canonical VFs as part of an immune evasion strategy (29). In chronically colonized patients, VF genes associated with acute infection may be under selective pressure to be lost or suppressed over time, resulting in a VF gene complement that is relatively stable across clinical states (29). This context may help explain why clinical state alone was insufficient to drive detectable differences in this cross-sectional metagenomic analysis.

Other plausible explanations exist for the non-significant clinical state finding. First, metagenomic DNA profiling captures gene presence and relative abundance but cannot distinguish between actively transcribed genes and silenced or dormant genetic material (30). Thus, sampling RNA rather than DNA may more accurately capture the transcriptionally active community of genes; the transcriptional activity of VF genes during exacerbation may differ meaningfully from gene abundance alone. Second, polymicrobial interactions may increase effective virulence without a concurrent change in individual gene abundance, as inter-species signaling between *P. aeruginosa* and other airway microbes has been shown to alter VF gene expression in vitro (31,32). Third, the small sample size of 22 pwCF contributing 75 respiratory samples for analysis limits statistical power, and it is possible that biologically meaningful differences exist but were not detectable in this cohort. Future studies with larger cohorts and metatranscriptomic designs will be needed to revisit this question more definitively.

### Pseudomonas aeruginosa Virulence Factors in High-Burden Samples

In contrast to the clinical state analysis, the comparison of high vs. low *P. aeruginosa* abundance samples identified biologically meaningful patterns in VF gene abundance. While there is no direct literature describing the role of aminotransferase degT as a *P. aeruginosa* VF in pwCF, other studies have shown aminotransferases to be important for pyoverdine production (33). Pyoverdine has been demonstrated as a key virulence determinant in *P. aeruginosa* isolates cultured from respiratory samples from pwCF (34) and is associated with iron acquisition (35).

Motility is recognized as a critical VF for *P. aeruginosa* and has long been established to adaptively decrease in the transition from acute to chronic and persistent infection (36–38). This is thought to be related to improved immune evasion, specifically through resistance to phagocytosis (39). Thus, the continued enrichment of fliQ, pilX, pilQ, and flgB in our dataset of CF samples with high *Pseudomonas* burden likely reflect active or recently established colonization rather than longstanding chronic persistence, which is line with our mostly adolescent study population.

The enrichment of T3SS, T4SS, and T6SS within our high-*Pseudomonas* samples is also notable. While the T3SS and T4SS inject bacterial proteins into eukaryotic cells, the T6SS works as a contact-dependent killing mechanism that *P. aeruginosa* can deploy against competing bacteria (40). Thus, the presence of T6SS-associated genes in high-burden samples may reflect the bacterium’s need to outcompete coinfecting organisms, a plausible strategy in the polymicrobial CF airway (41).

Although the enrichment of motility and secretion system genes in high-*Pseudomonas* samples aligns with expectations based on published virulence biology, it is important to note that these associations are correlational. Higher pathogen burden co-occurring with higher virulence gene abundance is not sufficient to establish a causal relationship between specific VFs and clinical outcomes in this dataset. It is equally possible that samples with high *P. aeruginosa* counts simply reflect greater total bacterial DNA, leading to proportionally more VF gene reads without a qualitative change in virulence capacity. Normalization strategies and longitudinal sampling would help address this interpretive limitation in future work.

### Staphylococcus aureus Virulence Factors and Antimicrobial Resistance

The *S. aureus* VF gene analysis yielded several findings that are both expected and clinically significant. The strong enrichment of Mph(C) family macrolide 2ʹ-phosphotransferase in high-*Staphylococcus* burden samples is consistent with the well-documented emergence of macrolide resistance in CF *S. aureus* isolates, likely driven by prophylactic use of azithromycin in this population. Phaff et al. demonstrated that long-term azithromycin therapy in children with CF is associated with the emergence of macrolide-resistant *S. aureus*, with resistance rates increasing from approximately 7% at baseline to over 53% within five years of therapy initiation (42). The presence of this resistance gene in high-burden samples in our cohort suggests that organisms with established macrolide resistance are more successful colonizers under the selective pressure of antibiotic prophylaxis, a finding that has direct clinical implications for therapeutic decision-making. The enrichment of tetracycline resistance protein in high-*Staphylococcus* samples, while less prominently discussed in the CF-specific literature (43), reflects a broader pattern of co-selection for multiple antibiotic resistance determinants observed in CF airways. *S. aureus* in CF environments is under substantial and multifaceted antibiotic selective pressure, and the accumulation of resistance genes across antibiotic classes is an expected evolutionary outcome. Nonetheless, the co-occurrence of macrolide and tetracycline resistance genes in high-burden samples underscores the concern that patients with higher *S. aureus* burdens may harbor organisms with more limited treatment options.

The enrichment of exotoxins and pathogenicity island proteins in high-*Staphylococcus* samples was among the most striking findings of this analysis. Staphylococcal pathogenicity islands (SaPIs) are mobile genetic elements encoding superantigen toxins, which are capable of activating large fractions of the T-cell repertoire and precipitating systemic immune responses disproportionate to bacterial burden (44). Fischer et al. reported that approximately 79.8% of *S. aureus* isolates cultured from the respiratory tract of persons CF at two geographically distinct centers harbored all six members of the enterotoxin gene cluster (EGC), suggesting that superantigen-encoding strains may be selectively maintained in the CF airway environment (45). Our data align with this observation: samples with higher *S. aureus* abundance showed enrichment of SaPI-associated genes, raising the possibility that superantigen-expressing strains are disproportionately represented among high-burden colonizers. The clinical implications of superantigen burden in CF airways are not fully understood, but chronic low-level superantigen exposure could contribute to the exuberant airway inflammation characteristic of CF lung disease.

The enrichment of vicinal oxygen chelate (VOC) family proteins in high-*Staphylococcus* samples, while less well characterized in the CF literature, is consistent with the broader theme of metabolic adaptation to the CF airway environment. VOC superfamily enzymes are involved in the detoxification of electrophilic compounds (46,47) and may contribute to the organism’s resilience in chemically hostile environments, including the oxidative and antibiotic-laden CF mucus layer. This finding is exploratory and requires further functional validation to determine its clinical significance.

Together, these results indicate that higher *S. aureus* abundance in CF airways is associated with enrichment of both antimicrobial resistance and virulence-encoding genes, including superantigen toxin carriers, which may have implications for disease severity and treatment response.

### Limitations

Several limitations of this study warrant explicit acknowledgment. First, and most critically, the small sample size (N = 75 respiratory samples) substantially limits statistical power. The failure to detect significant differences between clinical states may reflect a true null finding, but it may equally reflect insufficient power to detect effects of the magnitude present in this population. All significant findings from the pathogen abundance comparisons should be interpreted with corresponding caution, as multiple testing without pre-specified corrections inflates the risk of false positives.

Second, this study relied on metagenomic DNA sequencing, which captures gene presence and relative abundance but not expression. Gene abundance is a necessary but not sufficient condition for VF activity; active transcription, protein production, and post-translational regulation are all required for functional virulence. Future studies incorporating metatranscriptomics or metaproteomics would provide a more complete picture of virulence activity in the CF airway.

Third, the cohort had a mean age of 14.6 years at enrollment, limiting the generalizability of findings to younger children and older adults, in whom the microbial ecology of the CF airway can differ substantially. Similarly, the high prevalence of *P. aeruginosa* colonization (47%) suggests a cohort skewed toward later-stage or more severe disease in this mostly adolescent population, which may not represent the full spectrum of CF presentations encountered clinically.

Lastly, the use of relative abundance metrics in metagenomic studies is inherently compositional: an apparent increase in one organism’s abundance may partially reflect a decrease in others, rather than an absolute increase in pathogen burden. This compositional constraint should be considered when interpreting differential abundance findings.

### Future Directions

Several avenues of investigation emerge from these findings. The most immediate priority is replication in a larger, ideally multicenter cohort with sufficient power to detect clinically meaningful differences in virulence gene abundance between clinical states, age groups, and CFTR genotypes. The introduction of CF transmembrane conductance regulator (CFTR) modulators (e.g., elexacaftor/tezacaftor/ivacaftor) into standard CF care has substantially altered the airway microenvironment of many pwCF; future studies should account for modulator use as a key covariate, as these therapies may reduce pathogen burden and alter VF selection pressure.

The integration of metatranscriptomics with metagenomic sequencing would allow investigators to distinguish VF gene presence from active expression, addressing the fundamental interpretive limitation of DNA-only analyses. Paired DNA-RNA sampling from the same clinical specimens, with appropriate stabilization protocols, is technically feasible and would substantially strengthen causal inference in future studies.

The enrichment of antimicrobial resistance genes—particularly macrolide and tetracycline resistance—in high-*S. aureus* burden samples raises pragmatic clinical questions. Future work should examine whether VF gene and AMR gene co-enrichment is associated with measurable differences in clinical outcomes, including exacerbation frequency, hospitalization duration, lung function trajectory, or antibiotic treatment failure. Prospective studies linking metagenomic VF gene profiles to clinical outcomes would provide the evidence base necessary to translate these findings into actionable therapeutic guidance.

Finally, the role of inter-species dynamics, particularly the interaction between *P. aeruginosa* and *S. aureus*, in shaping the CF VF gene landscape deserves dedicated investigation. The observation that both pathogens were commonly co-detected in this cohort, combined with evidence that *P. aeruginosa* can deploy T6SS against *S. aureus*, suggests that pathogen competition may actively shape VF gene expression in vivo. Studies that explicitly examine the polymicrobial virulence interactome, rather than treating each pathogen in isolation, are likely to yield richer insights into CF lung infection biology.

## Conclusion

This study used shotgun metagenomic sequencing to characterize the VF gene landscape in the airways of 22 pwCF during pulmonary exacerbation. Contrary to our hypothesis, no statistically significant differences in VF gene abundance were observed between clinical states, a finding that may reflect the biological stability of the VF gene complement in chronically colonized CF airways rather than a failure of the pathogen to respond to clinical state. These results are consistent with emerging literature suggesting that exacerbations are not always accompanied by measurable increases in pathogen burden or VF gene expression in established chronic infections.

When samples were stratified by pathogen abundance, distinct VF gene signatures emerged. High *P. aeruginosa* burden was associated with enrichment of flagellar motility (FliQ) and type VI secretion system (TssG) genes, consistent with colonization competitiveness and host-pathogen interaction roles for these factors in the CF lung. High *S. aureus* burden was associated with enrichment of macrolide and tetracycline resistance genes, as well as pathogenicity island proteins encoding superantigen toxins, findings that align with the antibiotic selective pressure and virulence biology documented in prior CF-specific *S. aureus* literature.

These findings should be interpreted cautiously. The small sample size limits statistical power, the DNA-based design precludes conclusions about active VF gene expression, and the relative abundance framework introduces compositional constraints. Larger, longitudinal, and transcriptomic studies are needed to determine whether these VF gene patterns have measurable consequences for clinical outcomes and whether they could ultimately inform more targeted antimicrobial or anti-virulence therapeutic strategies in pwCF.

## Data Availability Statement

The sequence dataset supporting the conclusions of this article is available in the NCBI SRA repository under BioProject PRJNA825831.

## Author Contribution Statement

MLV performed bioinformatic and data analysis, generated the figures, and wrote the original manuscript. BH performed bioinformatic analysis. AB performed data collection, sample processing, and DNA extraction. AH conceptualized and designed the study, obtained funding, and supervised all aspects of the study, including selection of analysis methods, accuracy of analysis, and interpretation of findings. All authors were involved in critical review and editing of the manuscript and approve of the final version as written.

## Acknowledgements

The authors would like to acknowledge Dr. Claudio Anselmi and the Office of the Chief Research Information Officer for assistance and computing time on the high-performance computer at Children’s National Research and Innovation Campus (HPC@CNRIC).

## Generative AI Statement

During the preparation of this work, the authors used Claude to find manuscripts relevant to this topic and to provide suggestions to improve clarity of written content. After using these AI tools, the authors reviewed and edited the manuscript and take full responsibility for the content of the published article.

## Supplemental Results

**Supplemental Figure 1.**
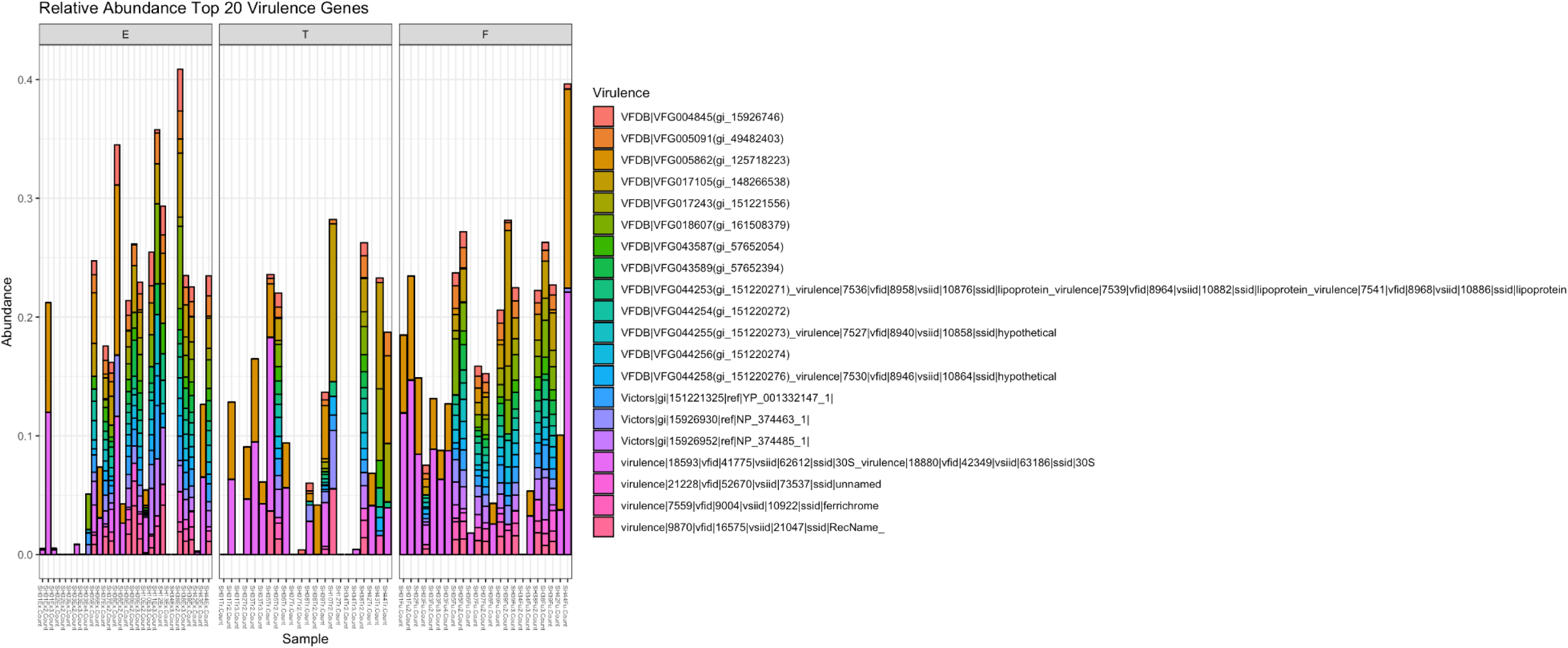
Relative abundance of the top 20 virulence factor genes, pulmonary exacerbation (E) versus completion of antibiotic treatment (T) versus follow-up (F).

**Supplemental Figure 2.**
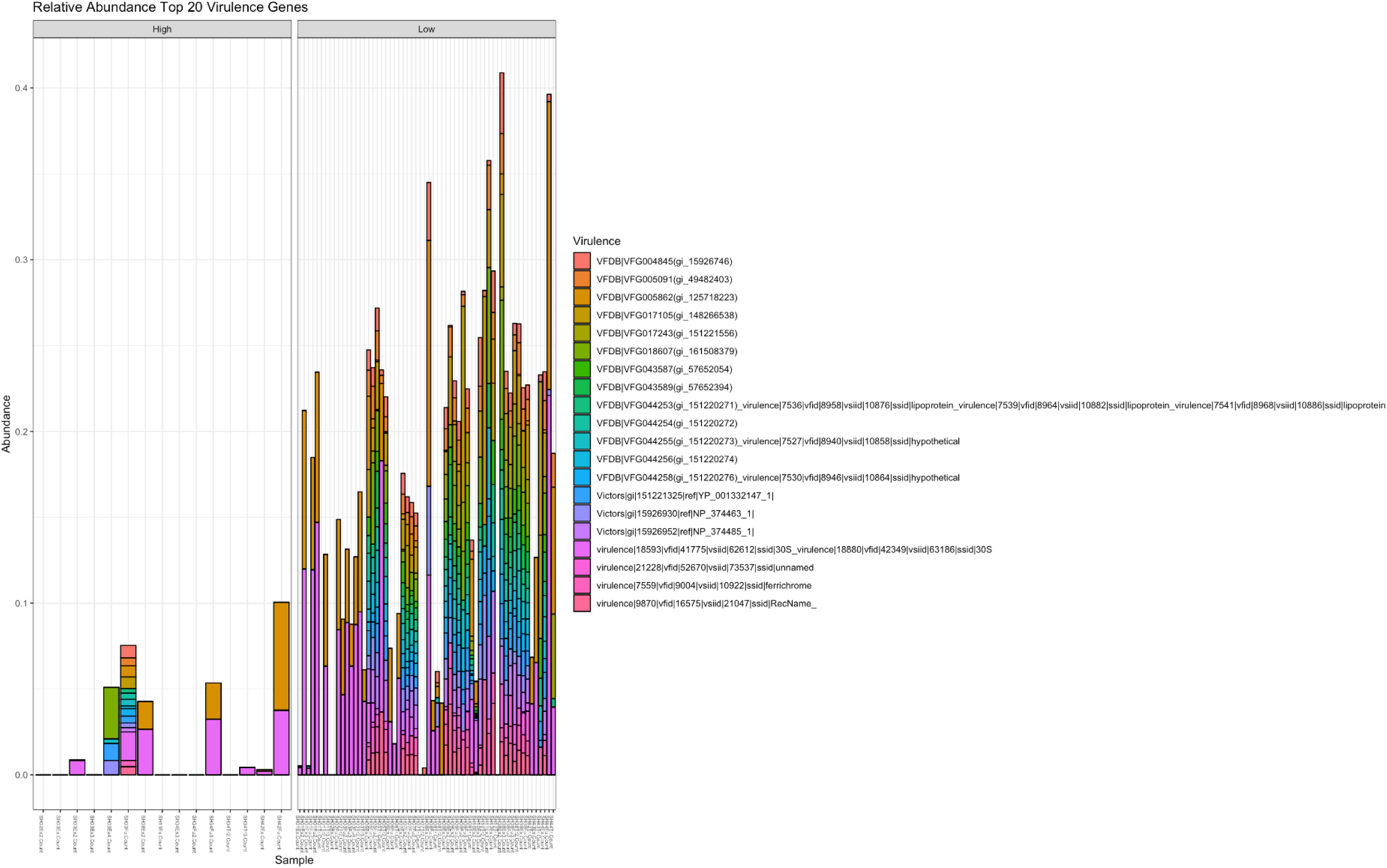
Relative abundance of the top 20 virulence factor genes, *Pseudomonas aeruginosa* high versus low.

**Supplemental Figure 3.**
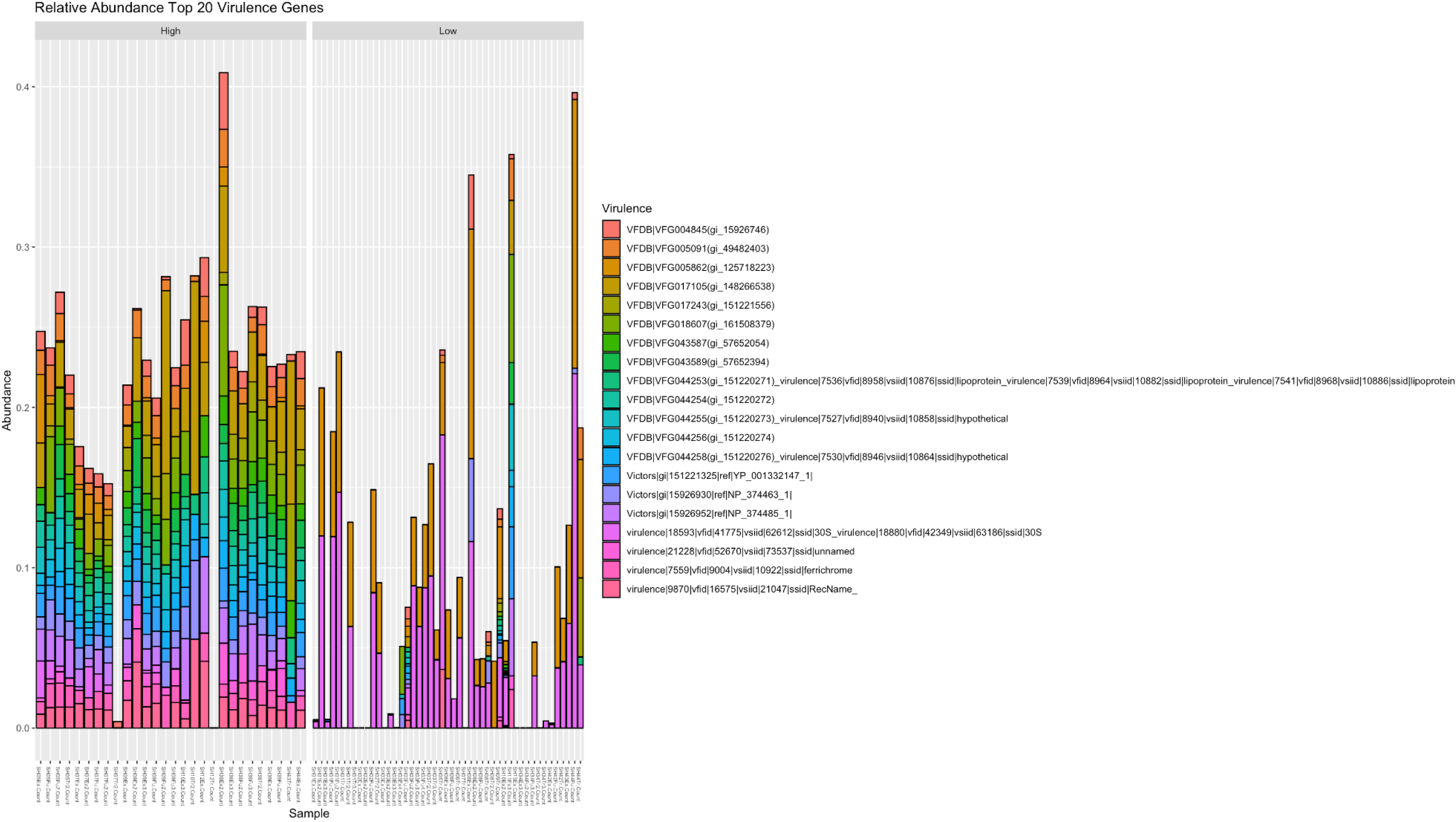
Relative abundance of the top 20 virulence factor genes, *Staphylococcus aureus* high versus low.

