## Supplementary figures and images for "Bacterial Virulence Genes Detected by Metagenomic Sequencing in the Cystic Fibrosis Airway Microbiome"

### Supplemental Figure 1

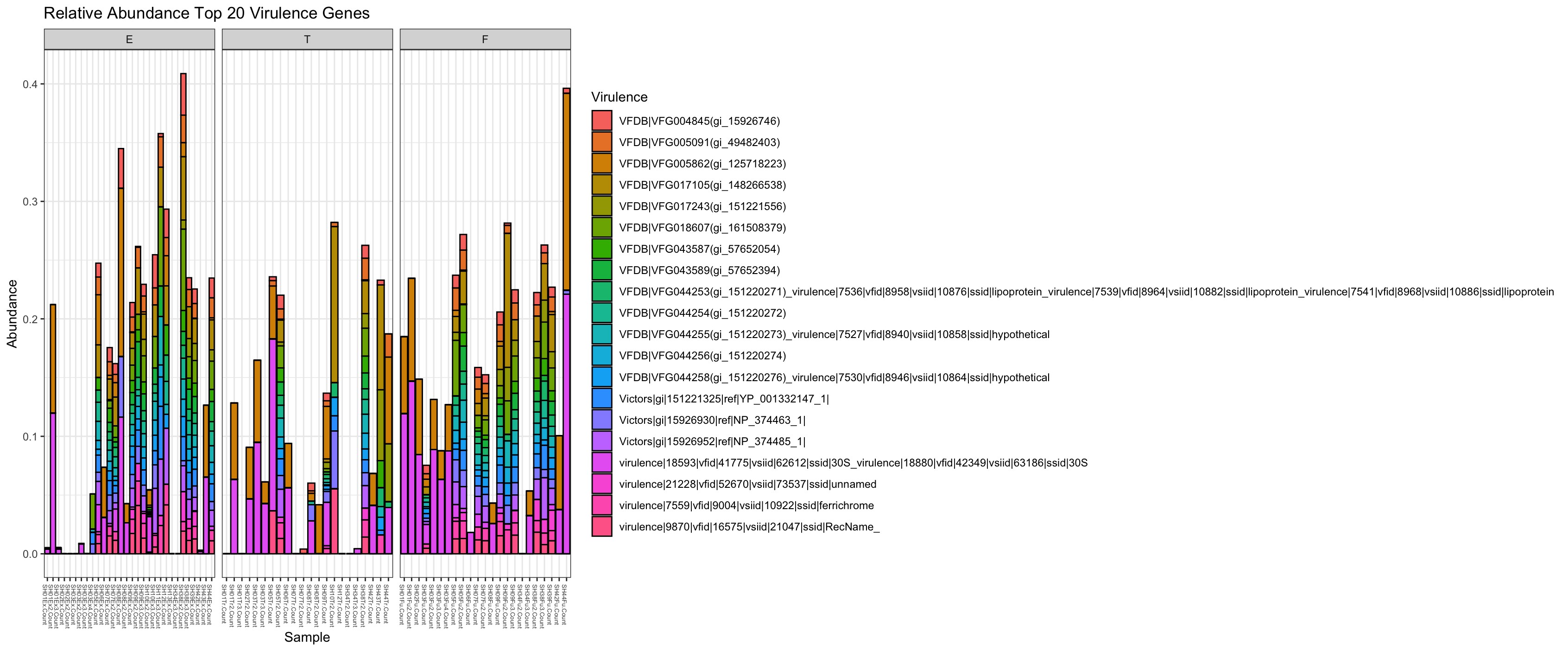

### Supplemental Figure 2

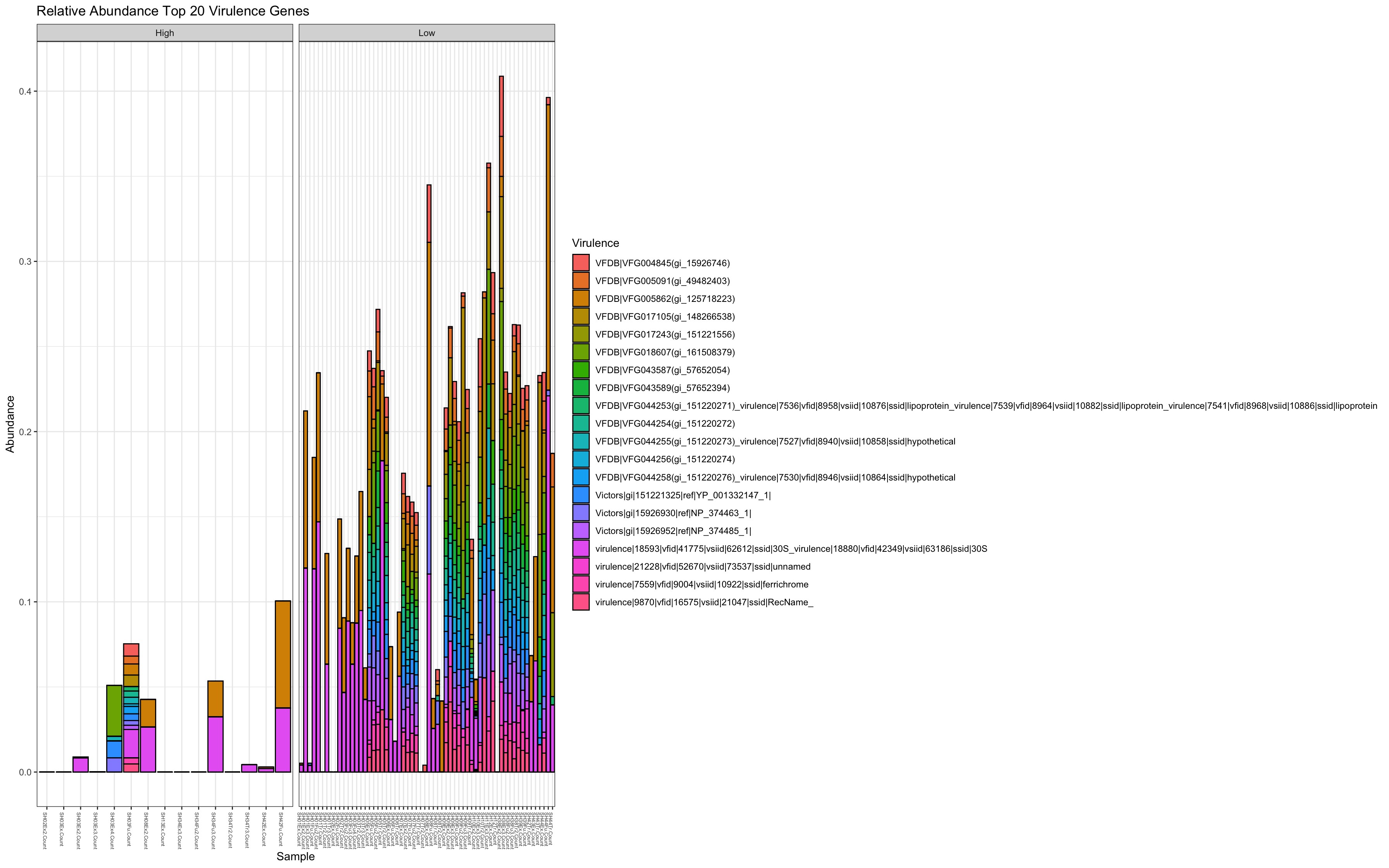

### Supplemental Figure 3

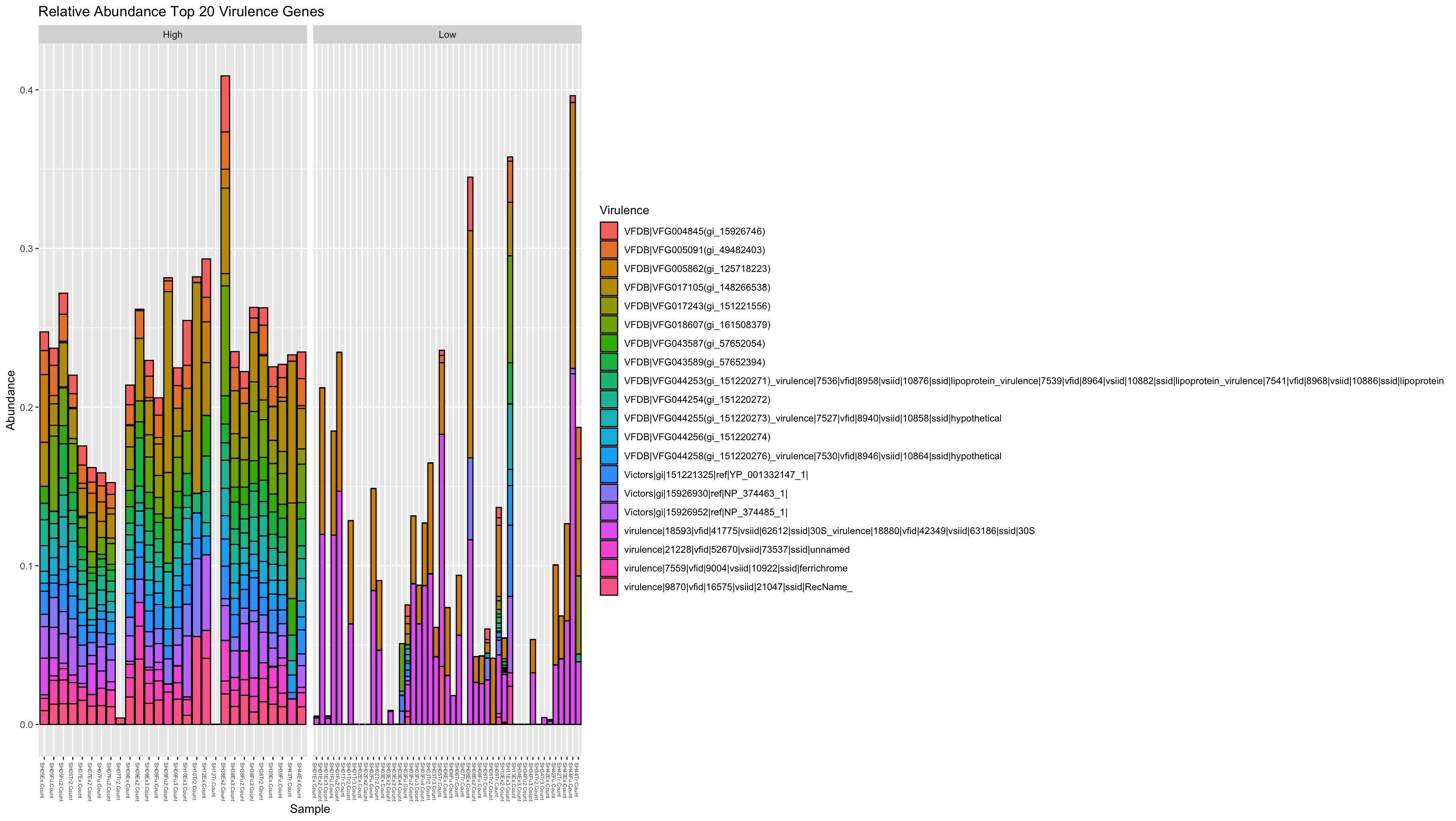
